# Tau Internalization is Regulated by 6-O Sulfation on Heparan Sulfate Proteoglycans (HSPGs)

**DOI:** 10.1101/167874

**Authors:** Jennifer N. Rauch, John J. Chen, Alexander W. Sorum, Gregory M. Miller, Tal Sharf, Stephanie K. See, Linda C. Hsieh-Wilson, Martin Kampmann, Kenneth S. Kosik

## Abstract

The misfolding and accumulation of tau protein into intracellular aggregates known as neurofibrillary tangles is a pathological hallmark of neurodegenerative diseases such as Alzheimer’s disease. However, while tau propagation is a known marker for disease progression, exactly how tau propagates from one cell to another and what mechanisms govern this spread are still unclear. Here, we report that cellular internalization of tau is regulated by quaternary structure and have developed a cellular assay to screen for genetic modulators of tau uptake. Using CRISPRi technology we have tested 3200 genes for their ability to regulate tau entry and identified enzymes in the heparan sulfate proteoglycan biosynthetic pathway as key regulators. We show that 6-O-sulfation is critical for tau-heparan sulfate interactions and that this modification regulates uptake in human central nervous system cell lines, iPS-derived neurons, and mouse organotypic brain slice culture. Together, these results suggest novel strategies to halt tau transmission.

---

The Microtubule-Associated Protein Tau (MAPT or tau) is an intrinsically disordered protein that under pathological conditions aggregates into filamentous inclusions known as neurofibrillary tangles (NFTs)^1^. While the composition and structure of NFTs are well characterized^2,3^, the *in vivo* process of aggregation is not well understood. The presence of NFTs is characteristic of a number of human diseases, collectively termed tauopathies. In tauopathies, such as Alzheimer’s disease (AD), NFT pathology advances in a predictable pattern throughout the brain affecting regions involved in learning and memory^4^. This progression of NFT pathology correlates with cognitive decline in patients and permits neuropathological diagnoses of patients in different stages of AD^5^.

The spread of protein aggregates during disease progression is a common theme in many neurodegenerative diseases, including α-synuclein in Parkinson’s disease^6^, Huntingtin protein in Huntington’s disease^7^, and superoxide dismutase-1 in amyotrophic lateral sclerosis^8^. However, the exact mechanisms underlying intercellular spread of these aggregates, including tau, is unclear. Increasing evidence suggests the transmission of tau pathology is mediated by the release, uptake, and trafficking of pathogenic or misfolded tau aggregates within synaptically connected neurons^9,10^. Once internalized, misfolded tau proteins act as a seed that recruits soluble endogenous tau into growing aggregates^11^. Aggregated tau is proteotoxic in model systems, suggesting that oligomeric and/or fibrillar tau may contribute to neurodegeneration^12^. However, it is unclear how the different quaternary architectures of tau affect internalization and if all structures have the ability to transfer between neurons. Further, a better understanding of the cellular processes that are necessary for transmission of tau aggregates could lead to the discovery of novel therapeutic strategies that would inhibit the spread of tau pathology and its consequences. Recent work on α-synuclein has shown that a cell surface receptor, lymphocyte activation gene-3 (LAG3), can bind α-synuclein and trigger its endocytosis into neurons^13^. We hypothesized that perhaps a receptor could also exist for tau.

Heparan sulfate proteoglycans (HSPGs) are a diverse family of proteins modified with the linear sulfated glycosaminoglycan (GAG) heparan sulfate (HS). HSPGs are present in virtually all cells and are involved in a multitude of processes including cell attachment, migration, differentiation, and inflammation^14^. HS chains consists of a basic disaccharide building block β1-4-linked D-glucuronic acid (GlcA) and α1-4-linked *N*-acetyl-D-glucosamine (GlcNAc). During assembly, HS chains can be highly modified; GlcNAc residues can be *N*-deacetylated and *N*-sulfated, GlcA can be epimerized at C5 to L-iduronic acid (IdoA), and ester-linked sulfate groups can be installed at C2 of the GlcA/IdoA and/or at C6/C3 of the GlcNAc. There is no defined template for HS modification on HSPGs. Thus, the availability of precursor material and abundance of biosynthetic enzymes in the cell are thought to dictate chain length, sulfation pattern, and epimerization^15^. Previous work has implicated HSPGs as a potential receptor for tau internalization^16,17^. However, this work has not examined whether specific HSPGs proteins or specific HS modifications on these proteins can act as determinants for tau entry, and therefore does not contain the molecular detail that would guide therapeutic strategies. Recent studies have also suggested that fragments of tau can discriminate between different sulfation modifications on heparin^18^. Within the cell there are a variety of enzymes known to be important for imparting this information on cellular receptors, thus we envisioned that there might be a specific protein or motif that would allow specification of tau entry.

In this work we used central nervous system (CNS) cell lines, iPS-derived neurons, and organotypic slice culture to understand the guidelines for tau uptake. We find that the quaternary structure of tau dictates the efficiency of uptake and we used tau monomers to screen for genetic modulators of internalization. We found that HSPG-modifying enzymes influence tau internalization and knockdown of these enzymes can repress the uptake of tau. We discovered that 6-O sulfation patterns on HS chains are critical for tau binding, and competition or removal of these motifs on the cell surface reduced the internalization of tau.

## Results

### Internalization of tau in CNS culture is regulated by quaternary structure

Tau protein can form multiple quaternary structures in solution, and recent evidence suggests that small tau oligomeric species may play a critical role in the spread of tau pathology and neurotoxicity^12^. Furthermore, work on smaller fragments of tau have suggested that the size of tau can regulate seeding capacity^17^. Therefore, to study the rules that govern the transmission of full length tau, we tested the uptake capacity of various tau structures in human CNS-derived cell lines and iPS-derived neurons.

First, we recombinantly expressed and purified the longest isoform of tau, 2N4R, and used established protocols to produce oligomeric and fibrillized tau species (see Methods). Characterization of these constructs using non-reducing SDS-PAGE analysis, revealed that tau monomers appear as one distinct band at ~64kDa, while tau oligomers/fibrils show additional bands of high molecular weight presumably for disulfide-bonded dimer, trimer, etc (Fig. 1a). TEM preparations of fibril samples displayed characteristic long helical filaments that could be fragmented upon sonication (Fig. 1a). We employed Dynamic Light Scattering (DLS) analysis to further characterize our tau constructs^19^. Analysis of our different tau species (Fig. 1b) showed, as expected, that monomeric tau was monodispersed and small in average size (5.7 ± 0.9nm), whereas oligomers, sonicated fibrils and fibrils were larger and more dispersed (19 ± 3nm, 33 ± 11nm, and 80 ± 8nm respectively).

**Figure 1.**
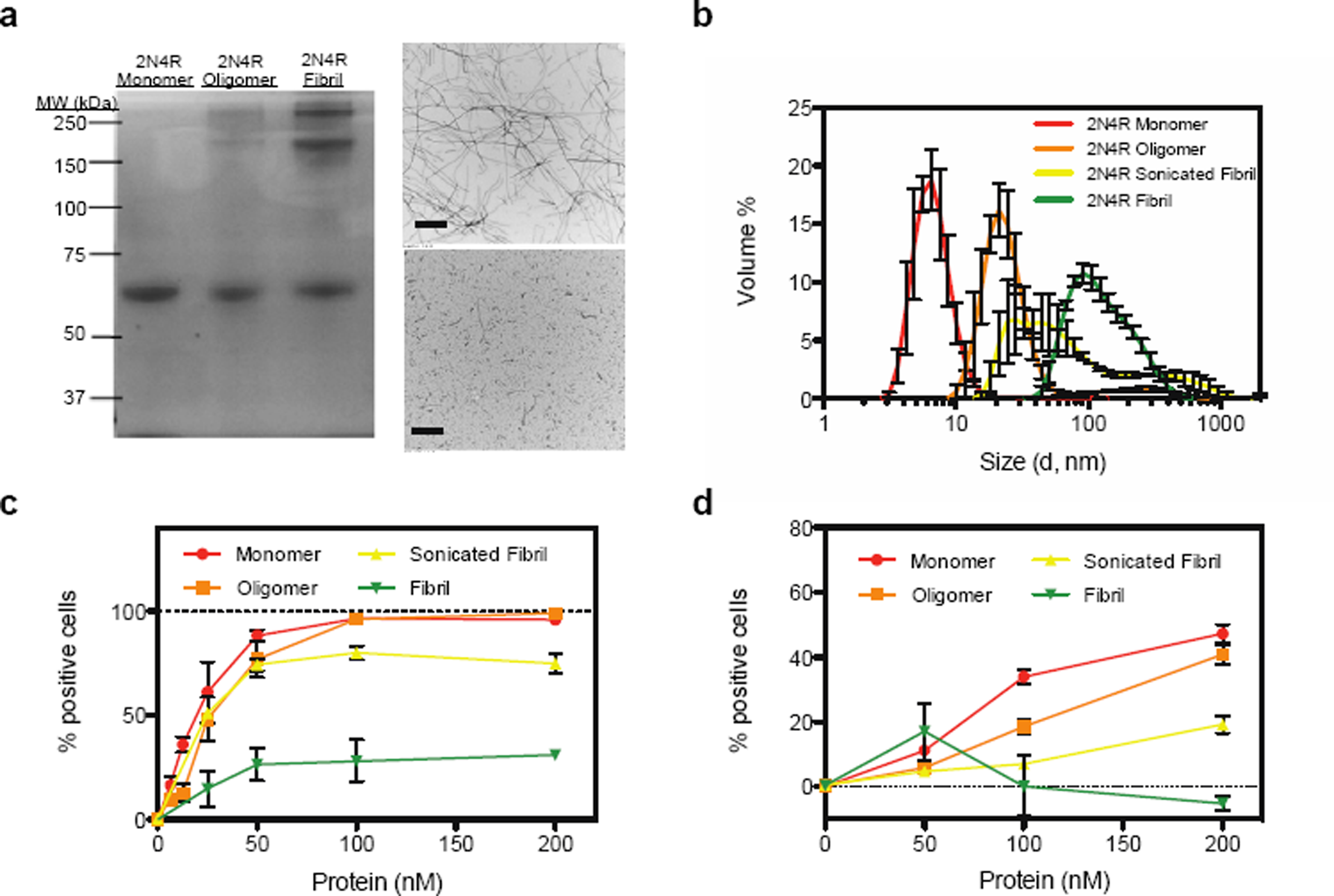
Tau uptake is regulated by quaternary structure. (**a**) 2N4R monomer, oligomer, and fibrillized proteins show distinct banding patterns on a non-reducing SDS-PAGE gel. 2N4R Fibrils before (top) and after (bottom) sonication as visualized by negative-stain EM. Bar represents 500nm (**b**) DLS of 2N4R monomer, oligomer, fibril, and sonicated fibril species. Experiments were performed in triplicate and the error shown is SD. (**c**) Uptake of 2N4R quaternary structures in H4 cells. (**d**) Uptake of 2N4R quaternary structures in iPS-derived neurons. For all uptake experiments, three independent experiments were performed in duplicate, identical control experiments were performed at 4°C and subtracted from 37°C data to generate final curves shown, and the error shown is SEM.

To test the efficacy of these tau species towards internalization by cells, we labeled each protein preparation with an Alexa Fluor-488 (AF488) probe and then added various concentrations of protein to the cell media of H4 neuroglioma cells. After one hour, cells were washed, lifted from the plate and analyzed for fluorescence using flow cytometry. To control for non-specific binding to the membrane, identical experiments were performed at 4°C (a non-permissive temperature for endocytosis), and any fluorescence observed was subtracted from our results. Monomeric, oligomeric and sonicated fibrils were efficiently internalized, while fibril samples were not (Fig. 1c). Tau uptake was a time-dependent process, with uptake observed in as little as 10 minutes (Supplementary Fig. 1a). This assay for tau uptake was robust for other human CNS cell lines, including SHSY-5Y neuroblastoma, and ReN VM neural progenitors (Supplementary Fig. 1b,c). Further, human iPS-derived neurons also showed a preference for smaller structures of tau, with fibrillized tau showing nearly no uptake (Fig. 1d). Taken together, these results suggest that cellular uptake of tau is regulated by size and that large species of tau are inefficiently internalized across multiple cell types.

### Functional genomics to find modulators of tau uptake

With a robust and high-throughput assay for tau uptake in hand, we used this platform to screen for genetic modulators that could either increase or decrease tau internalization. To do this, we developed an H4 CRISPRi cell line that stably expresses a catalytically inactive Cas9 fusion protein (dCas9-KRAB). CRISPRi represses transcription of genes with high specificity using single guide RNAs (sgRNAs) that guide the dCas9-KRAB protein to the transcription start site (TSS) of the targeted gene^20^. As a proof of concept, we screened a next-generation CRISPRi library of sgRNAs that targeted 3200 different genes with five different sgRNAs per gene, and contained hundreds of non-targeting negative-control sgRNAs^21^. H4 CRISPRi cells transduced with sgRNAs were then incubated with 25nM AF488 labeled 2N4R monomer (2N4R-488) for 1hr and sorted based on AF488 fluorescence. The cell populations with the top and bottom thirds of AF488 fluorescence, representing higher than average and lower than average tau uptake, respectively, were recovered, genomic DNA was isolated and the locus encoding the sgRNA was PCR-amplified and frequencies of each sgRNA in the two populations were determined by next-generation sequencing (Fig. 2a). To detect hit genes, we applied our previously developed quantitative framework for pooled genetic screens^22,23^ as described in the Methods.

**Figure 2.**
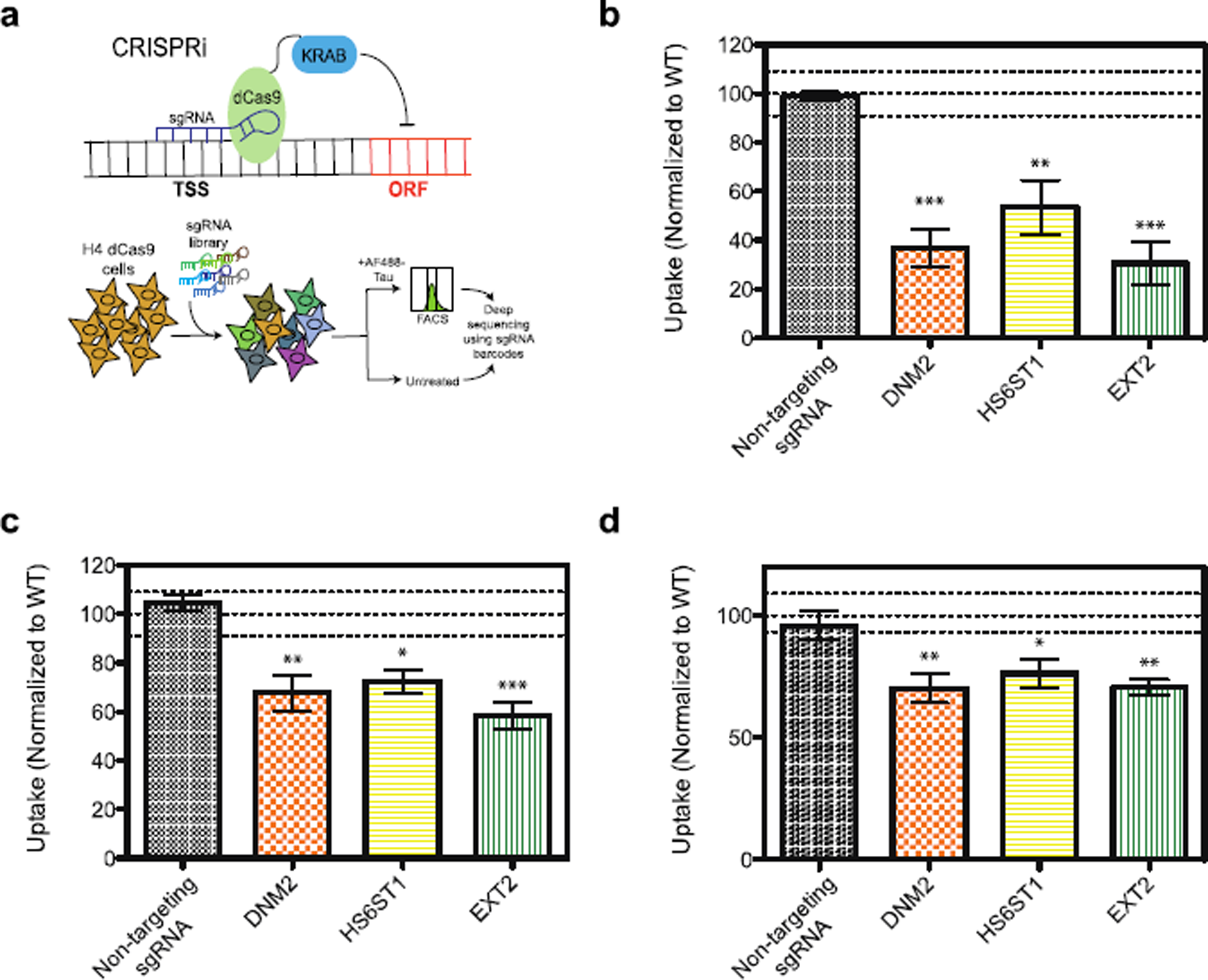
CRISPRi screen for tau uptake modulators. (**a**) Screening strategy using H4 CRISPRi cells, see text for details. (**b**) Reconfirmation of selected hits in H4i cells with 2N4R monomer (50nM), normalized to a WT (no sgRNA) control. (**c**) Uptake of 2N4R oligomers (50nM) in H4i cells with selected gene knockdowns, normalized to a WT (no sgRNA) control. (**d**) Uptake of 2N4R monomer in iPS-derived neurons with selected gene knockdowns, normalized to a WT (no sgRNA) control. All uptake experiments were performed in duplicate over three independent experiments with the data combined. Lines on the graphs represent WT mean +/− 3 standard deviations, error bars represent SEM, a one-way ANOVA analysis with Dunnett’s method was used to determine significance between the gene knockdown and the non-targeting sgRNA *=p-value ≤0.05, **=p-value ≤ 0.01, ***=p-value ≤0.001.

We selected sixteen genes for individual validation studies, including genes with particularly strong phenotypes in the primary screen, as well as genes in pathways previously implicated in tau uptake. Fifteen of the sixteen follow-up hits were found to repress expression of their target gene (i.e. knockdown) as determined by qPCR (Supplementary Fig. 2a). Fourteen of the fifteen gene knockdowns also reproduced their screen phenotype i.e. either increased or decreased tau uptake as compared to a non-targeting sgRNA control (Fig. 2b and Supplementary Fig. 2b).

Interestingly, we found that some of the strongest phenotypes in our screen were attributed to genes that are known cell cycle regulators (Supplementary Fig. 2c). The selective knockdown of genes such as TP53, led to a decrease in G1 length and thus an overall increase in cell proliferation (Supplementary Fig. 2d). Endocytosis is known to increase during G1 phase^24^; therefore, it seemed logical that cell cycle regulators that can shorten the G1 phase could reduce the amount of tau uptake and vice versa. In line with these observations a small molecule inhibitor of CDK4/6 that causes a stall in G1 phase (PD0332991) was sufficient to almost double the amount of tau taken up in H4 cells (Supplementary Fig. 2e). Further work will be needed to dissect if and how these genes might influence post-mitotic neurons.

Hits that were of particular interest included genes involved in heparan sulfate proteoglycan (HSPG) biosynthesis (EXT2 and HS6ST1) as well as DNM2, a GTPase involved in endocytosis. These single gene knockdowns repressed uptake of tau monomer by over 50% (Fig. 2b) and also reduced the uptake of tau oligomers (Fig. 2c). Further, these gene knockdowns were sufficient to reduce the uptake of tau in iPS-derived neurons (Fig. 2d). Treatment of cells with an inhibitor for DNM2, Dynasore, was also able to reduce uptake of tau (Supplementary Fig. 2f).

### Tau binds to heparin derivatives and shows specificity for 6-O-sulfated heparins

Based on our pilot screen results, we were particularly interested in the enzymes involved in HSPG biosynthesis. Previous reports had indicated that internalization of tau could be regulated by HSPGs^16^, but our identification of HS6ST1, an enzyme that is responsible for 6-O sulfation of HSPGs, supports a hypothesis that specific motifs on HSPGs might be important for tau uptake.

HSPGs are decorated with HS chains that consist of a repeating GlcA-GlcNAc building block. During assembly, HS chains can be highly modified and, importantly for their function, they can be sulfated at multiple sites within the disaccharide (Fig. 3a). To ascertain whether specific sulfation motifs were important for tau binding, we employed a heparin ELISA assay. In this assay, biotin-labeled heparin derivatives (Fig. 3b) were immobilized on streptavidin-coated plates, purified tau protein was incubated with the plate at increasing concentrations, and antibodies were used to detect tau binding. Tau bound heparin with an affinity of 4.6 ± 0.6nM, consistent with previous reports^25^. Over sulfated heparin (>3 sulfates per disaccharide) bound tau even tighter (2.2 ± 0.5nM), while fully-desulfated heparin showed a drastically reduced binding (>1μM) (Fig. 3c). Removal of *N*-sulfates or 2-O sulfates had little effect on tau binding (15.0 ± 5nM and 7.4 ± 1.0nM, respectively), while removal of all O-sulfates and, in particular 6-O sulfates, led to a significant decrease in tau binding (>1μM in each case; Fig. 3c,d). The relative binding of 6-O desulfated heparin in the presence of 10nM tau was 6.9% that of heparin, consistent with our hypothesis that tau interacts with the 6-O sulfation motif on HS chains, and that HS is important for tau internalization.

**Figure 3.**
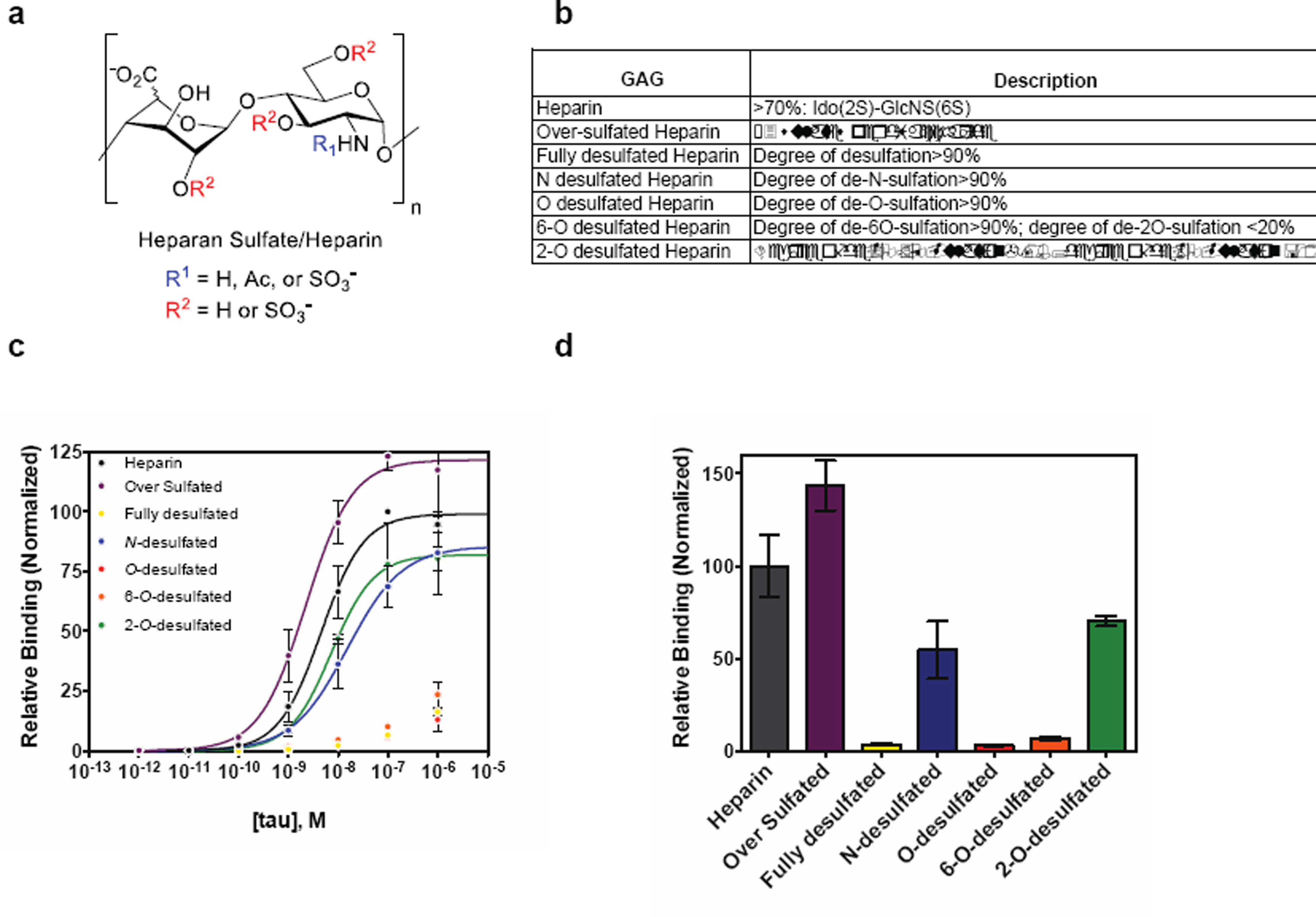
Binding of tau to heparin derivatives. (**a**) HS chains consist of GlcA/IdoA-GlcNAc disaccharide units that can be modified at positions indicated in red and blue. (**b**) List of heparin/heparin derivatives that were used and description of their sulfation modifications. (**c**) Binding of 2N4R to various heparin derivatives by ELISA with data fit to a Hill binding model (line) where appropriate. (**d**) Normalized relative binding of 2N4R (10nM) to various heparin derivatives. Three independent experiments were performed, data were normalized to heparin controls, and the error shown is SD.

### Cellular internalization of tau is effected by the presence of 6-O-sulfation

To confirm that 6-O sulfation was indeed a determinant of tau uptake, we developed a cellular competition experiment to monitor tau internalization. In this experiment, we added heparin or heparin derivatives free into the cell media immediately prior to addition of 2N4R-488. We found that internalization of tau can be efficiently competed by the presence of heparin or HS in the media (Fig. 4a). Likewise, addition of 2-O desulfated heparin was able to reduce uptake, whereas 6-O desulfated heparin or chondroitin sulfate (negative control) were significantly less effective at reducing uptake (Fig. 4a). These results were consistent when tested in iPS-derived neurons (Fig. 4b), demonstrating that the 6-O sulfation motif is indeed a critical determinant for cellular tau entry.

**Figure 4.**
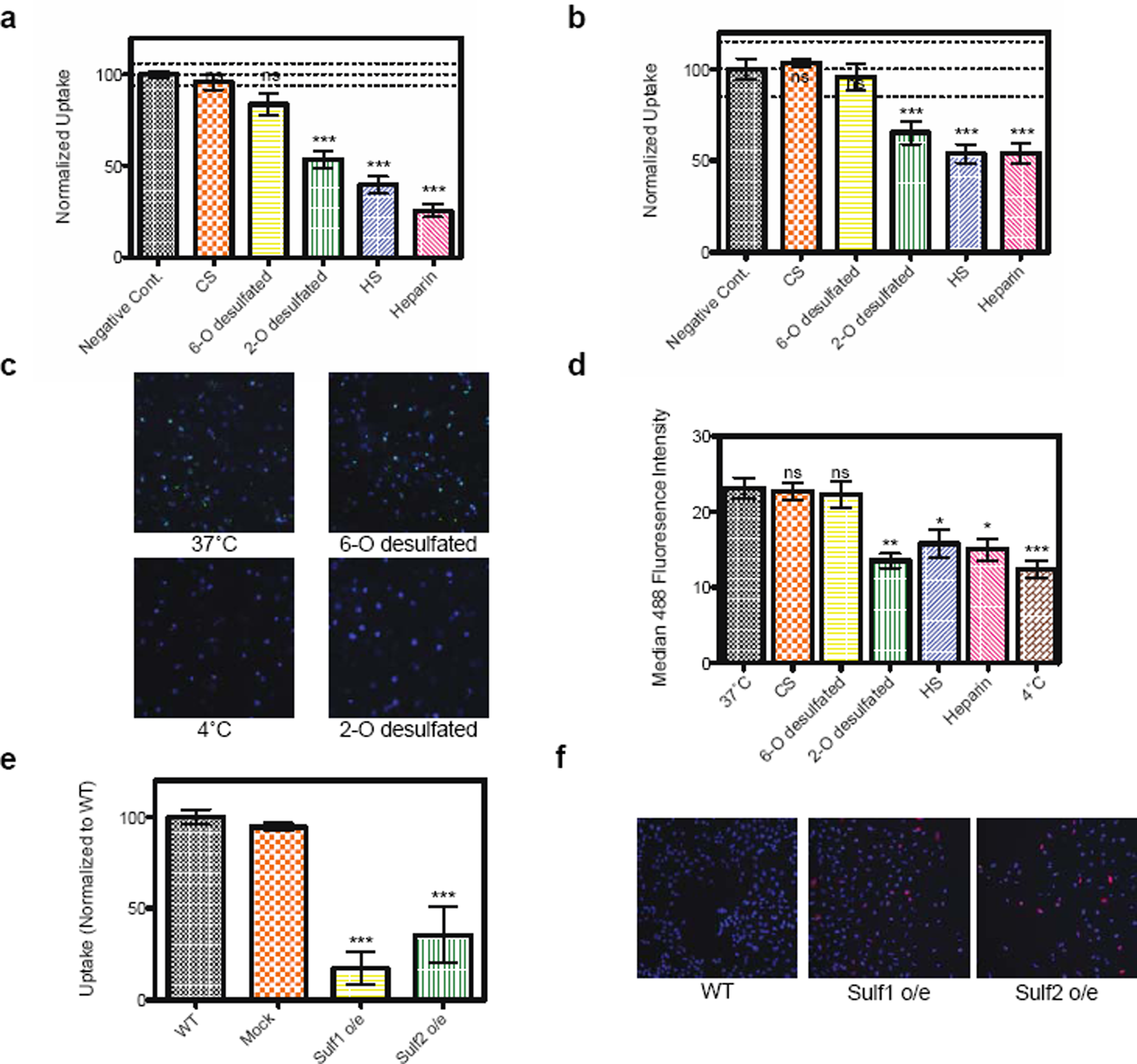
6-O sulfation regulates tau uptake in CNS culture. (**a**) Internalization of 2N4R-488 into H4 cells is strongly inhibited by incubation with heparin, heparan sulfate, and 2-O desulfated heparin as compared to 6-O desulfated heparin or chondroitin sulfate. (**b**) Internalization of 2N4R-488 into iPS-derived neurons is inhibited by incubation with heparin, heparan sulfate, and 2-O desulfated heparin as compared to 6-O desulfated heparin or chondroitin sulfate. All uptake experiments were performed in duplicate over three independent experiments with the data combined. Lines on the graphs represent WT mean +/− 3 standard deviations, error bars represent SEM, and a one-way ANOVA analysis with Dunnett’s method was used to determine significance compared to the negative control *=p-value ≤ 0.05, **=p-value ≤0.01, ***=p-value ≤0.001, ns = not significant. (**c**) Internalization of 2N4R-488 into mouse organotypic slice culture is strongly inhibited by incubation with 2-O desulfated heparin, but not 6-O desulfated heparin. Hoechst stain is used to label nuclei. (**d**) Quantification of the median 488 fluorescence intensity for tau uptake in mouse slice culture. Two independent experiments were performed with multiple images (>5) from each condition. Error bars represent SEM. (**e**) Uptake of 2N4R-488 tau is reduced when Sulf1 or Sulf2 is overexpressed in H4 cells as compared to WT or mock transfected cells. Uptake experiments were performed in duplicate over three independent experiments with the data normalized to WT and combined. Error bars represent SEM. (**f**) ICC confirms that Sulf1 and Sulf2 are overexpressed (red) in H4 cells. Hoechst stain is used to label nuclei.

To test if HS 6-O sulfation would also prevent internalization of tau in an *ex vivo* system, we generated organotypic brain slices from adult mice using established methods^26^. These slices showed normal electrophysiology, suggesting good vitality (Supplementary Fig. 3a). Their ability to uptake tau was tested by incubating the cultures with 2N4R-488 for 30min at 37°C or, as a control, 4°C. The cultures were washed, stained with Hoechst dye to label nuclei, and mounted on coverslips to image. Incubation at 37°C showed uptake of tau in the slice cultures, whereas very little fluorescence was observed in the 4°C control cultures (Fig. 4c and Supplementary Fig. 3b). Consistent with our previous results, incubation with heparin, heparan sulfate, or 2-O desulfated heparin reduced uptake of tau as quantified by the median 488 fluorescence intensity (Fig. 4d). Chondroitin sulfate and 6-O desulfated heparin incubation did not reduce the median fluorescence, verifying that 6-O sulfation is also important for tau internalization *ex vivo* (Fig. 4d and Supplementary Fig. 3b). Quantification of median Hoechst fluorescence across all the images showed that similar cell numbers were analyzed for each condition (Supplementary Fig. 3c).

Finally, in order to test if direct removal of 6-O sulfates from the cell surface could reduce uptake of tau in cell culture, we overexpressed two different extracellular endosulfatases (Sulf1 & Sulf2) that selectively cleave 6-O sulfates on GlcNAc^27^. Overexpression of these enzymes in H4 cells, showed a dramatic decrease in tau uptake with Sulf1 reducing uptake to 17 ± 9%, and Sulf2 reducing uptake to 36 ± 15% (Fig. 4e). Overexpression of the constructs was confirmed with immunocytochemistry (Fig. 4f).

## Discussion

In recent years, a growing body of literature has reinforced the hypothesis that prion-like cell-to-cell propagation of protein aggregates underlies disease progression of numerous neurodegenerative disorders^28^. For some of these protein aggregates (PrP, α-synuclein), specific pathways and protein conformers that are key for transmission have been delineated quite fully^13,29^. However for other proteins, such as tau, many details regarding its transmissibility are still unclear. It has been shown in various cellular and animal models that exogenously added tau aggregates can induce tau pathology^9,30^; however, the disparity between the experimental systems and the tau species used in each study makes it hard to draw a unifying theme. Tau is a complex protein that has several isoforms, posttranslational modifications, and the ability to form multiple quaternary structures^31^. Therefore, we have focused our research on the development of a cellular assay that can robustly detect internalization across various tau constructs and CNS cell systems.

The data presented here demonstrate that full-length human tau (2N4R) can be efficiently internalized across multiple relevant cell systems and that this uptake is dependent on the overall size of the tau species. This is a critical point, since it is still debated exactly how tau induces neurotoxicity. One hypothesis suggests that tau oligomers are the toxic species and that the formation of tau fibrils inside neurons (NFTs) is a protective mechanism. This is supported by neuropathological studies in AD patients where neuronal loss and cognitive deficits precede NFT formation^32^. Our data are consistent with this hypothesis, as tau fibrils were unable to be internalized, and thus would be unable to “seed” misfolding inside the cell. This does not, however, discount the potential for synaptic transmission or exosome transfer of fibrils between neurons^33^. But, as tau oligomers and monomers have been found elevated in the CSF of AD patients^34,35^, it seems likely our assay recapitulates at least one *in vivo* scenario.

Using CRISPRi technology, we identified multiple genetic factors that can influence the uptake of tau in cell culture. The identification of HSPGs as regulators of tau internalization is not an entirely new concept. Indeed, it has been known for years that tau can bind heparin, and that this molecule can be used as an *in vitro* inducer of aggregation36. However, our work has shown that binding of tau to HS is directly related to the sulfation pattern, and specifically on the presence of 6-O sulfates on the glucosamine subunit. Further work will be needed to understand if there are specific HS chain lengths or regions on tau that are important for this interaction.

This work is an important step forward in the characterization of HS-tau interactions. With these new insights we can begin to envision ways to design small molecules that mimic the HS structures that interact with tau to block its aggregation and neutralize its toxicity. It is plausible to expect that novel treatments that target the HS-tau interaction may contribute to AD treatment and could improve the effects of other treatments.

## Methods

### Chemicals

Heparin, chondroitin sulfate, heparan sulfate, and desulfated heparins (Neoparin & Galen Labs Supplies). Dynasore hydrate (Sigma). Hoechst (Thermo Scientific).

### Protein Purification, Labeling and Fibrillization

Full length tau protein (2N4R, 1-441aa) was purified with slight modification to previously published protocols^37^. Briefly, 2N4R tau in the pRK172 plasmid was expressed in *E. coli* BL21 (DE3). Cell pellets were harvested and resuspended in cell lysis buffer (50mM MES pH 6.5, 5mM DTT, 1mM PMSF, 1mM EGTA) + cOmplete protease inhibitor tablets (Roche). Lysate was sonicated, boiled for 10min, and then centrifuged at 50,000xg for 30min at 4°C. The supernatant was then precipitated with ammonium sulfate (20% w/v) and centrifuged at 20,000xg for 30min at 4°C. The pellet was resuspended into 4mL of MonoS Buffer A (50mM MES pH 6.5, 50mM NaCl, 2mM DTT, 1mM PMSF, 1mM EGTA) and dialyzed overnight against the same buffer. The protein was loaded onto a MonoS column (GE Healthcare) and eluted with a linear gradient of NaCl using MonoS Buffer B (Buffer A + 1M NaCl). Fractions containing 2N4R tau were pooled, concentrated, and dialyzed overnight into PBS pH 7.4. Protein concentration was determined using a BCA assay (Thermo Scientific).

Protein was labeled with Alexa Fluor^®^ 488 or 647 5-SDP ester (Life Technologies) according to the suppliers instructions. After labeling, 100mM glycine was added to quench the reaction and the proteins were subjected to Zeba desalting columns (Thermo Scientific) to remove any unreacted label. Average label incorporation was between 1 and 1.5 moles/mole of protein, as determined by measuring fluorescence and protein concentration (A_max_ x MW of protein / [protein] x ε_dye_). To prepare tau fibrils and oligomers, 10uM protein, in PBS 1mM DTT pH 7.4, was mixed with heparin (0.05mg/ml) and incubated with shaking at 37°C. The formation of oligomers was observed after 4h of shaking, whereas fibril formation was formed after 5 days^30^. To make sonicated fibril samples, fibrillized protein was sonicated using a MiSonix Sonicator 4000 (QSonica, LLC) at 50% amplitude for 60 1s pulses.

### Transmission Electron Microscopy

Tau fibrils and sonicated fibrils (1μM) were absorbed on 200-mesh formvar-coated copper grids, washed, and stained with a 2% uranyl acetate solution. Grids were then imaged with a JEOL JEM-1230 (JEOL USA, Inc) at the indicated magnifications.

### Dynamic Light Scattering

Protein solutions (1μM) were filtered (0.45μm) and analyzed using a Zetasizer Nano ZS (Malvern). The time-dependent autocorrelation function of the photocurrent was acquired every 10s, with 15-20 acquisitions for each run and with at least three repetitions. The error bars displayed on the DLS graphs were obtained by the standard deviation (SD) between replicates.

### Cell Culture, Transfections and Treatments

H4 cells were cultured in DMEM supplemented with 10% FBS, 100μg/ml penicillin/streptomycin. SHSY-5Y cells were cultured in DMEM/F12 supplemented with 10% FBS, 100μg/ml penicillin/streptomycin. ReN-VM cells were cultured on Matrigel (Corning) coated plates in DMEM/F12 supplemented with 2μg/ml heparin (STEMCELL Technologies), 2% B27 (Life Technologies, 100μg/ml penicillin/streptomycin, 20μg/ml bFGF (Stemgent), 20μg/ml EGF (Sigma). Cultures were maintained in a humidified atmosphere of 5% CO_2_ at 37°C. Transfection of H4 cells with Sulf1 (Addgene #13003) or Sulf2 (Addgene #13004) were performed with Lipofectamine 3000 (Invitrogen) according to the manufacturers instructions and cells were assayed 48h later. Overexpression was confirmed with immunocytochemistry using the his tag epitope (see supplemental methods). For competition experiments, heparin (and various derivatives) was added to the media just prior to tau addition at the indicated concentrations.

### iPS Culturing and Differentiation

CRISPRi iPSc with an inducible TRE3G-dCas9KRAB-T2A-mCherry were maintained in 6-well Matrigel (Corning) coated plates with mTeSR1 media (STEMCELL Technologies) and split with ReLeSR (STEMCELL Technologies) at a 1:20 ratio every 4-5 days. For differentiation, CRISPRi cells were split with Accutase and virally infected twice with a NeuroD1-IRES-eGFP-Puro (Addgene #45567) according to previously published methods^38^. Doxycyclin was added to induce expression of NeuroD1, and the following day cells were selected with puromycin (5μg/ml). After selection, cells were lifted with Accutase treatment and replated 1:12 in PEI coated 24-well plates. Three days later, media was changed to Brain Phys Neuronal Medium (STEMCELL Technologies) with doxycycline and AraC to remove any remaining dividing cells. At day 8, cells were infected with specific sgRNA Technologies, 100μg/ml penicillin/streptomycin, 20μg/ml bFGF (Stemgent), 20μ g/ml μg/ml EGF (Sigma). constructs. iPS neurons were assayed between days 14-18 of maturity as described below. Immunocytochemistry was used to confirm neuronal properties (Supplementary Fig. 1d).

### Flow Cytometry

H4 cells were plated at 50,000 cells per well in a 24-well plate. The next day media was replaced, and cells were treated with varying concentrations of AF488-labeled tau protein for 1hr (unless indicated otherwise) at 37°C. Cells were then washed twice with PBS and trypsinized to lift cells from the plate. Identical control experiments were performed at 4°C to confirm that tau protein was internalized and not just adhering to the cell membrane. Lifted cells were analyzed using an Accuri-C6 Flow Cytometer and propidium iodide was used to exclude dead cells from the analysis.

### CRISPRi Screen

We generated a stable CRISPRi-enabled H4 neuroglioma cell line by transducing H4 cells with the lentiviral plasmid pHR-SFFV-dCas9-BFP-KRAB^39^ and selecting a polyclonal population of BFP-expressing cells by FACS. These cells were transduced with a next-generation CRISPRi sgRNA library (sublibrary “Cancer and Apoptosis”)^21^, and transduced cells were selected using puromycin (1μg/mL). Media was replaced with fresh DMEM containing AF488-labeled Tau monomers at a final concentration of 25nM for 1 hr at 37°C. Cells were washed twice, trypsinized, and resuspended in FACS buffer (PBS with 0.5% FBS). A BD-FACAria II, was used to sort live cells into two populations based on the top and bottom thirds of AF488 fluorescence; approximately 15 million and 17 million cells were recovered from the high- and low-fluorescence populations, respectively. The resulting DNA was isolated, the cassette encoding the sgRNA was amplified by PCR, and relative sgRNA abundance was determined by next-generation sequencing as previously described^20,23^. We analyzed the resulting data using our previously developed quantitative framework for pooled genetic screens^22,23^. Statistical significance for each targeted transcription start site was calculated using the Mann-Whitney U test to compare the phenotypes of the 5 sgRNAs targeting the transcription start site to the phenotype distribution of the 280 non-targeting sgRNAs in the library.

## ELISA

Heparin/heparin derivatives were biotinylated using EZ-link Sulfo-NHS-LC-Biotin (Thermo) and immobilized on streptavidin-coated plates, followed by a 2 hr incubation with 2N4R tau protein containing a C-terminal myc tag (10-fold dilution series). Bound tau was detected using an anti-myc HRP-conjugated antibody (Bethyl, A190-105P), visualized using TMB substrate (R&D Systems), and quantified by UV-Vis. Absorbance for each plate was measured at 450 and 550nm. The 550nm measurement is a correction for plate imperfections and was subtracted from the 450nm values. Data were normalized to heparin controls and fit with a Hill binding mathematical model where appropriate using the equation *Y=B_max_*X^H*/(*K_d_^H + X^H)*, where H is the Hill slope (variable), X is the concentration of tau, and B_max_ is the binding maximum.

### Organotypic Slice Culture & EPSP measurements

Slice cultures were prepared as previously described^26^. Hippocampal excitatory postsynaptic potentials (EPSPs) were measured using a multi-electrode array (MEA). Stimulus current was injected into a region of the hippocampal slice by an MEA electrode, with the magnitude of stimulus current ranging from 20-80 μA. EPSPs were then measured by recording an electrode in a different region of the hippocampal slice. Smaller stimulus currents excite fewer cells, and thus the measured EPSPs have smaller magnitudes.

### qPCR

Purelink RNA Extraction Kit (Invitrogen) was used to isolate RNA from samples. RNA (1μg) was then converted to cDNA using SuperScript Reverse Transcriptase III (Invitrogen) according to the supplier’s instructions. Real-time quantitative PCR was performed using Power SYBR Green PCR Master Mix (Applied Biosystems) according to QuantStudioTM 12K Flex Real-Time PCR System protocol. GAPDH mRNA level was used to normalize samples.

### Immunocytochemistry

Cells were fixed with 4% paraformaldehyde for 15min at RT followed by three washes with PBS. Cells were permeabilized with 0.25% Triton X-100 in PBS, and blocked for 1hr in blocking buffer at RT (1% BSA, 300mM Glycine, 0.1% Gelatin, 4% Donkey Serum in TBST). After blocking, cells were incubated with primary antibodies diluted in blocking buffer overnight at 4°C. The day after cells were washed three times (5 min each) with 0.05% Tween-20 in PBS. The following primary antibodies were used: Tau-46 (Invitrogen, 36400, 1:1000), MAP2 (Millipore, AB5622, 1:1000), and anti-his (Thermo Scientific, MA1-21315, 1:1000). Secondary antibodies (Life Technologies, A21422, A21206, 1:1000) were incubated for 1hr at RT, washed three times with 0.05% Tween-20 in PBS and imaged with an Olympus IX71 Microscope

## Acknowledgements

We thank Jason Gestwicki (UCSF) for the 2N4R pRK172 plasmid, Bruce Conklin (UCSF) for the CRISPRi iPS cells, the UCSB NRI-MCDB Microscopy Facility for use of the TEM, the UCSB Stem Cell Core for use of the facility, and the UCSB BNL for access to the DLS.

J.J.C. was supported by a postdoctoral fellowship from the Alzheimer’s Association and the QB3/Calico Longevity Fellowship. S.K.S. was supported by a National Defense Science & Engineering Graduate (NDSEG) Fellowship. A.W.S. was supported by an NIH Training Grant (NIH/NRSA 5T32 GM007616-38). L.C.H.-W. was supported by an NIH/NIGMS grant (R01 GM093627). Support also came from an NIH Director’s New Innovator Award (NIH/NIGMS DP2 GM119139) (M.K.), an Allen Distinguished Investigator Award (Paul G. Allen Family Foundation) (M.K.), the Tau Center Without Walls (NIH/NINDS U54 NS100717) (M.K., K.S.K), the Tau Consortium (K.S.K.) the Chan-Zuckerberg Biohub (M.K.) and the Paul F. Glenn Center for Aging Research (M.K.).

## Author Contributions

J.N.R., J.J.C., A.W.S., G.M.M, and T.S. performed experiments and interpreted results. S.K.S. made and validated the H4 CRISPRi cell line. L.C.H.-W., M.K., and K.S.K., interpreted results and directed the research. J.N.R., J.J.C., L.C.H.-W., M.K., and K.S.K. wrote the manuscript.

